# Epigenetic Divergence during Early Stages of Speciation in an African Crater Lake Cichlid Fish

**DOI:** 10.1101/2021.07.30.435319

**Authors:** Grégoire Vernaz, Alan G. Hudson, M. Emília Santos, Bettina Fischer, Madeleine Carruthers, Asilatu H. Shechonge, Nestory P. Gabagambi, Alexandra M. Tyers, Benjamin P. Ngatunga, Milan Malinsky, Richard Durbin, George F. Turner, Martin J. Genner, Eric A. Miska

## Abstract

Epigenetic variation can alter transcription and promote phenotypic divergence between populations facing different environmental challenges. Here we assess the epigenetic basis of diversification during the early stages of speciation. We focus on the extent and functional relevance of DNA methylome divergence between two *Astatotilapia calliptera* ecomorphs in crater Lake Masoko, southern Tanzania. We report extensive genome-wide methylome divergence between populations linked to key biological processes, including transcriptional activity of ecologically-relevant genes. These include genes involved in steroid metabolism, haemoglobin composition and erythropoiesis, consistent with divergent habitat occupancy of the ecomorphs. Using a common garden experiment, we found that global methylation profiles are rapidly remodelled across generations, but ecomorph-specific differences can be inherited. Collectively, our study suggests an epigenetic contribution to early stages of vertebrate speciation.

**One sentence summary:** Inheritance and plasticity of epigenetic divergence characterise early stages of speciation in an incipient cichlid species of an African crater lake.

## Main text

The genomic basis of adaptive phenotypic diversification and speciation has been extensively studied but many questions remain (*1*–*3*). Recent studies in plants and animals have provided initial evidence for a contribution of heritable epigenetic divergence to functional phenotypic traits (*4*–*9*). However, whether epigenetic processes facilitate adaptive diversification, especially during the early stages of speciation, remains unknown.

To investigate the potential role of epigenetic processes during vertebrate speciation, we focus on the incipient *Astatotilapia* radiation in crater Lake Masoko, southern Tanzania, which is within the Lake Malawi catchment (Fig. 1A and Fig. S1A-C) (*10*). The two *Astatotilapia* ecomorphs present are characterised by different depth preferences, morphology, diet and male nuptial coloration (*10, 11*). Specifically, the littoral ecomorph (yellow males) occupies the well-oxygenated shallow waters (≤5m), while the benthic ecomorph (blue males) thrives in deeper (20-30m), almost anoxic and dimly lit habitats of the lake (Fig. 1B-C and Fig. S1C)(*12*). Previous genome-level sequencing has revealed overall low sequence divergence (*F*_ST_ = 0.038) between the two ecomorph pairs (*11*), consistent with early stages of divergence. However, there was elevated genetic differentiation (F_ST_>0.3) in 98 genomic regions, which were enriched for functional targets of divergent selection, notably in relation to vision, morphogenesis and hormone signalling (*2, 11*). Collectively, the two ecomorphs are monophyletic with respect to *Astatotilapia* in the neighbouring Mbaka river system (Fig. S1A), which they separated from ca. 10,000 years ago. More recently the colonisation and adaptation to the benthic habitat from littoral fish has taken place (1,000 years ago; 200-350 generations)(*11*). Against this background, the incipient *Astatotilapia* radiation of Lake Masoko provides a valuable system to investigate the role of epigenetic processes during the early stages of sympatric speciation. Here we combined reduced-representation bisulphite sequencing (RRBS), whole-genome bisulphite sequencing (WGBS) and whole transcriptome sequencing (RNAseq) to assess functional methylome divergence in the two *Astatotilapia* ecomorphs of Lake Masoko and their neighbouring ancestral riverine population.

**Fig. 1.**
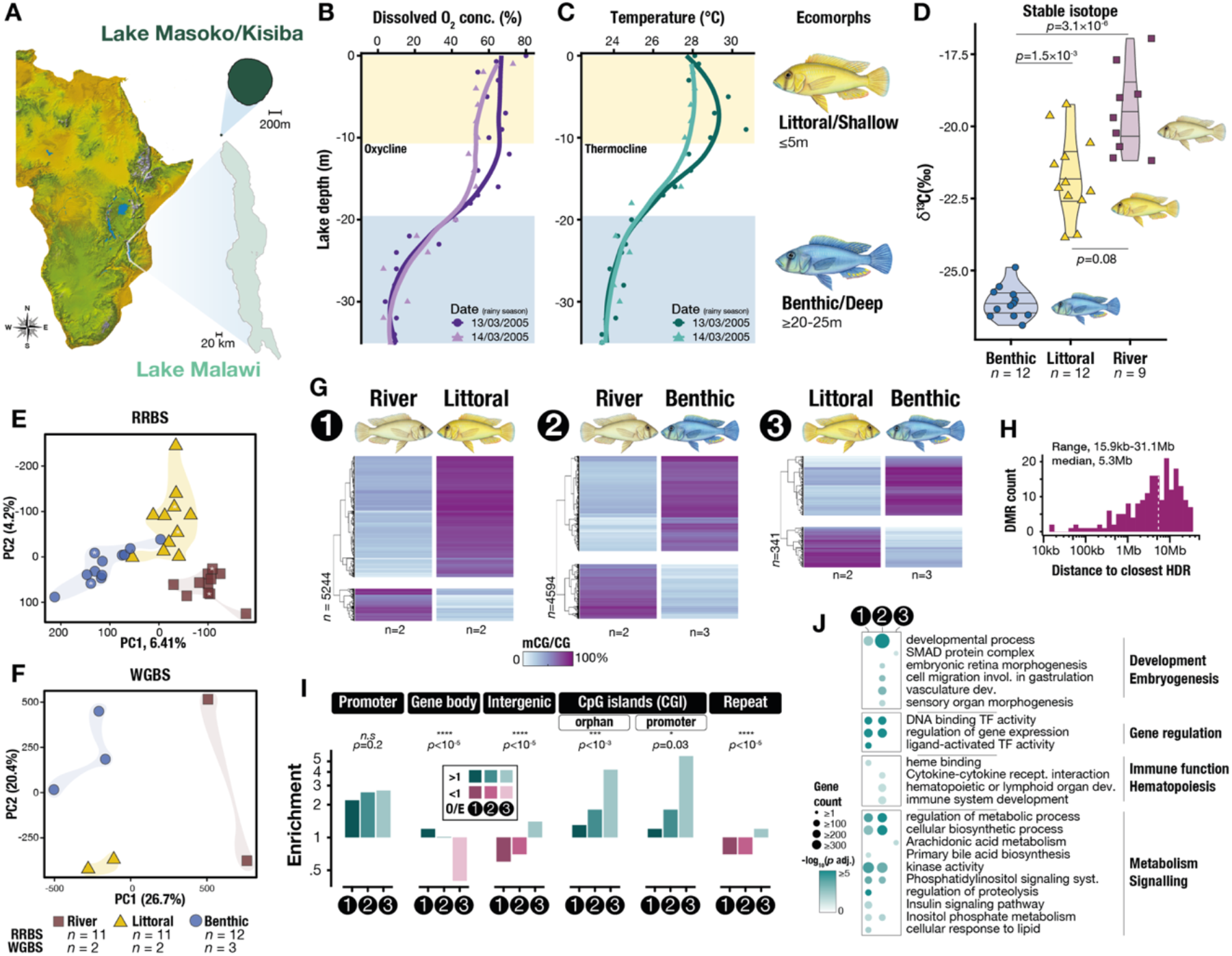
Whole-genome DNA methylation landscape of the *Astatotilapia* cichlid radiation in Lake Masoko. **A.** Geographical map of Lake Masoko (also known as Lake Kisiba), Tanzania (satellite image: NASA). **B**. Dissolved oxygen (O_2_) concentration (%) by depth (meters, m) in Lake Masoko. **C**. Water temperature (ºC) by depth. The oxycline (B) and thermocline (C) divide the habitats of the two *Astatotilapia calliptera* ecomorphs of Lake Masoko: the littoral (yellow) morph population thrives in shallow (≤5m), well oxygenated waters, while the benthic (blue) morph population is found in deeper, colder and almost anoxic habitats of the lake. Data from Ref.(*12*), collected on two different days at same depths (LOESS smoothed curves). **D**. Violin plots of stable isotope ratios (δ^13^C_V-PDB_, ‰) by population indicate a more offshore zooplankton-based diet for the benthic fish. *p*-values for Dunn’s multiple comparison tests are shown. *n* represents the number of fish per population. **E to F**. Principal component analysis (PC1-2) of liver methylome (mCG) variation using both the reduced representation bisulfite sequencing (RRBS)(E) and whole-genome bisulfite sequencing (WGBS)(F) datasets. PC1 and PC2 segregate the three populations apart in both datasets. Percentage of total variance given in parentheses. Asterisks in (E) indicate samples used for WGBS. *n* indicates the number of biological replicates for each population. **G**. Unbiased hierarchical clustering and heatmap of the average DNA methylation levels at all significantly differentially methylated regions found between the three pairwise comparisons (numbered 1-3) reveal population-specific methylome patterns. mCG/CG levels (%) averaged by population over each DMR (p≤0.05). Total number of DMRs for each comparison shown vertically. **H**. Histogram of closest distance in bp (log scale; median, dotted line) between DMRs and HDRs (when on same chromosome). **I**. Enrichment plots (observed/expected ratio) for methylome variation (DMRs) in different genomic features for each pairwise comparison. Chi-square tests (p values shown) were performed for the three population distributions for each genomic feature. Repeat: transposon-only repeats. **J**. Gene ontology enrichment analysis for genes associated with methylome differences (either in promoter, gene body or 1-5kbp away from a given gene) for each pairwise comparison.

### Population-specific methylome divergence during early stages of speciation

To examine genome-wide population-level methylome divergence, we combined two bisulphite sequencing approaches. First, we generated reduced-representation bisulphite sequencing (RRBS) data from liver tissue of 12 wild-caught adult males for each of the littoral and benthic *Astatotilapia* ecomorphs of Lake Masoko, and of 11 wild-caught adult males from the neighbouring Mbaka river. In addition, we produced whole-genome bisulphite sequencing (WGBS) data from liver tissue of at least two individuals from each population. On average, 11.1±3.4 million single-end 75bp-long reads and 341±84.3 million paired-end 150bp-long reads were generated per individual for the RRBS and WGBS dataset, respectively (Tables S1-S2; Methods). Reads yielded similar mapping rates to the SNP-corrected *Maylandia zebra* reference assembly across ecomorphs (GCF_000238955.4; see Methods), consistent with low inter-population sequence divergence (Fig. S2A-B; see Materials and Methods)(*11*). Principal component analysis (PCA) of both RRBS and WGBS datasets revealed strong methylome segregation among the three populations (Fig. 1E-F).

As functional methylation variation tends to occur over neighbouring CG dinucleotide sites, we can identify differentially methylated regions (DMRs; ≥25% methylation difference, ≥4 CpG, ≥50bp-long, *p* <0.05; Materials and Methods) between each pair of populations using the WGBS dataset. In total, 5244 DMRs were found between the riverine and littoral fish, and 4595 between riverine and benthic fish (Fig. 1G and Fig. S3A). The majority of these DMRs (79% and 63.5% total DMRs, respectively) showed substantial gain in methylation (gain-DMRs; ≥44% methylation increase) in the littoral or benthic fish compared to ancestral riverine fish, respectively, with average methylation levels at gain-DMRs of ≥72% mCG/CG (Fig.1G and Fig. S3 B-C). Between the littoral and benthic fish, 341 DMRs were found, two-thirds of which showed an increase in methylation in the benthic fish population. In all pairwise comparisons, DMRs varied in length from 50 bp to 3 kbp (median, 250 bp) and were distributed across all chromosomes (Fig. S3 D-E). To investigate the relationship between underlying genetic polymorphism and methylome divergence, regions of elevated genetic differentiation (highly diverged region [FST≥0.3], HDR; Ref. (*11*)) between the littoral and benthic populations were examined. These regions showed high methylome conservation and were not co-localised with DMRs. Notably, the distances between HDRs and the nearest DMRs ranged from 15.9 kb to 31.1 Mb (median, 5.3 Mb), suggesting that large regions of high genetic differentiation were in general not associated with epigenetic divergence between the two incipient Masoko ecomorphs (Figs. 1H and S3 G).

We next examined the genomic localisation of DMRs among populations, and found promoter regions to be highly enriched in all comparisons (∼2.5-fold enrichment over chance; one sample t-test *p*<0.0001), consistent with their *cis*-regulatory functions (Fig. 1I) (*13*). In the recent littoral to benthic transition, the differences in methylomes were particularly overrepresented in CpG-dense regions (CpG-islands; ≥4.5-fold enrichment; *p*<0.0001) located both within (pCGI) and outside promoters (orphan CGI/oCGI,). oCGIs may represent distant *cis*-regulatory regions, such as ectopic promoters or enhancers (*14*), both of which are known targets of methyl-sensitive DNA-binding proteins (MDP) (*13, 15*). In contrast methylome variation within intergenic regions and transposable elements only showed slight enrichment in the comparison of the Masoko ecomorphs relative to comparisons involving the riverine population. Additionally, the methylome divergence in gene bodies, the function of which remains unclear in vertebrates but could be involved in alternative splicing (*13*), was generally very low, and even highly underrepresented (2.5-fold depletion, *p*<0.0001) in the comparison of Masoko ecomorphs. This suggests high methylome conservation between gene bodies of the most closely related populations.

We identified biological processes associated with methylome divergence by performing gene ontology enrichment analysis (Fig. 1J). We first noted that genes involved in transcription regulation were highly enriched in DMRs, and then identified three further biological processes for the genes enriched in methylome divergence: immune function and haematopoiesis; embryogenesis and development; and metabolism (Fig.1J and Fig. S4). Notably, methylome divergence in developmental genes has been shown previously to account for close to half of all species-specific epigenetic differences between three species of the Lake Malawi cichlid radiation (*7*). Additionally, regions showing benthic-specific methylome patterns and located outside gene body were significantly enriched for specific transcription factor (TF) binding motifs (Fig. S3F), including the haematopoiesis-related SCL/Tal1 (stem cell leukaemia/T-cell acute lymphocytic leukaemia 1; *p*=1×10^−145^), forkhead hepatocyte nuclear factor 3-alpha (foxa1; *p*=1×10^−9^), the embryogenesis-related smad2 (mothers against decapentaplegic homolog 2; *p*=1×10^−125^) and the metabolic/circadian clock transcriptional activator bmal1 (aryl hydrocarbon receptor nuclear translocator-like protein 1; *p*=1×10^−8^), consistent with altered transcription factor activity. This suggests *cis*-regulatory functions for such population-specific DMRs in development, haematopoiesis and metabolism, associated with the adaptation to the benthic habitat.

### Methylome divergence associated with transcriptional changes in haematopoiesis and liver-specific metabolism

We investigated the link between population-specific methylome divergence and transcriptional activity using total RNA sequencing data from liver tissues of 4-5 individuals of each group (mean ± sd, 32.9 ± 3.8 million paired-end 100/150bp-long reads per fish; Table S3). As with whole methylome variation, we observe population-specific transcription patterns (Fig. 2A). We then performed differential gene expression analysis (false discovery rate [FDR] adjusted p-value using Benjamini-Hochberg < 1%, fold change ≥1.5 and high gene expression in ≥1 population [top 90^th^ percentile]; see Materials and Methods) and found a total of 546 significantly differentially expressed genes (DEGs) between the three populations, including 125 genes that were differentially expressed between the Masoko ecomorphs (Fig. 2B). Close to 38% of all DEGs show reduced transcriptional activity in Masoko ecomorphs relative to the riverine population, and these genes were significantly enriched for functions related to energy balance/homeostasis and steroid metabolism, including the PPAR (peroxisome proliferator-activated receptor) and foxO (forkhead box O) signalling pathways, consistent with metabolic adaptation to different diets (Fig.2B). Conversely, almost all the remaining DEGs (52%) show high transcriptional activity almost exclusively in the benthic fish and were primarily enriched for functions associated with haemoglobin complex/oxygen binding activities and iron homeostasis as well as fatty acid metabolism, in line with the occupation of a hypoxic environment, and possibly related to their zooplankton-rich diet (Fig.1 A, D). Critically, significant changes in transcriptional activity were strongly associated with methylome divergence at promoter regions (44% of all DEGs, overrepresentation factor=5.3, p<10^−28^; Fig. 2C).

**Fig. 2.**
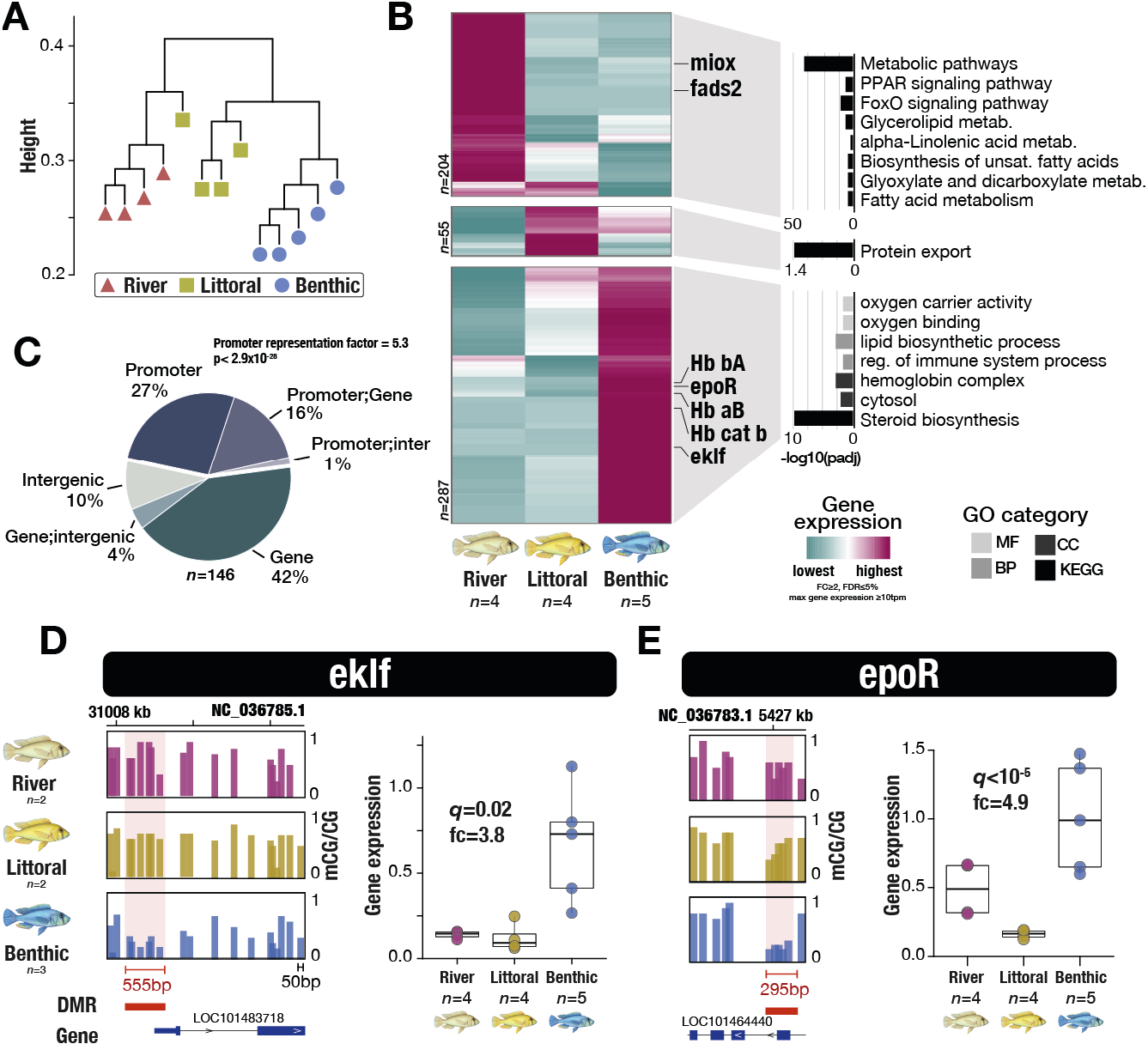
Methylome variation is associated with altered transcriptional activity. **A.** Unbiased hierarchical dendrogram based on whole transcriptome variation (Euclidean distances plotted). **B**. Unbiased hierarchical clustering and heatmap of transcriptional activity (*Z*-score) for all significantly differentially expressed genes (DEGs) among the three wild populations, showing three different clusters of transcriptional activity (false discovery rate adjusted p-value, using Benjamini-Hochberg [FDR] <0.05, fold change ≥2 and high gene expression [top 90^th^ percentile] in any one sample). Examples of differentially expressed genes are shown on the right-hand side of the graph. Right: gene ontology enrichment for DEGs of each of the three transcriptional activity clusters (only qval<0.01 shown). **C**. Pie chart representing the genomic localisation of DMRs associated with DEGs. Significant overlap between DMRs at promoter and transcriptional changes (representation factor= 5.3, p<10^−28^; exact hypergeometric probability). DEGs can be associated with multiple DMRs in different locations (promoter, intergenic and genic DMRs). **D-E**. The promoters of the genes erythroid transcription factor krüppel-like factor 1 (eklf) and the erythropoietin receptor (epoR), both involved in erythropoiesis and red blood cells differentiation, show hypomethylation levels in the livers of the benthic fish compared to the littoral populations. Genome browser view of methylome profiles for each ecomorph are shown. Each bar represents the average mCG/CG levels in 50bp-long non-overlapping windows for each ecomorph group. DMRs (p<0.05) are highlighted in red and the length of each DMR is indicated in red. Right-hand side of D and E: boxplots of gene expression in liver of benthic, littoral and river fish for eklf and epoR are shown (qval, FDR; fc, fold change).

Focusing on the functional categories identified above, we examined in detail several examples of transcriptional diversification associated with divergence in methylome landscapes. First, loss of methylation in two genes associated with active haematopoiesis and erythrocyte differentiation, the erythroid krüppel-like transcription factor (ekfl) and the erythropoietin receptor (epoR), was associated with significant gain of transcriptional activity in the benthic fish compared to the littoral population (fold changes ≥3.8, FDR<0.02; Fig. 2D-E). eklf plays an essential role in erythropoiesis, haem synthesis and in the modulation of globin genes expression (*16*), and its transcriptional activity has been linked to methylation changes previously in humans (*17*). Moreover, three haemoglobin subunit (Hb) genes, adjacent to each other within the MN (major) globin cluster on chromosome LG4 (NC_036783.1) were significantly upregulated (≥4.5-fold) in the benthic fish specifically (FDR<0.02; Figs. S5A). This includes the cathodic Hb beta (cHbB) which is known to have higher oxygen-binding affinity in anoxic environments and is expressed exclusively in benthic fish (*18*). Notably, a 2kbp-long DMR adjacent to this globin gene cluster shows benthic-specific hypermethylation methylome divergence compared to intermediate and unmethylated levels in littoral and river fishes respectively (Fig. S6). This suggests it bears *cis*-regulatory functions, possibly similar to the vertebrate Locus Control Regions (LCR), known to be bound by many essential erythroid transcription factors, such as eklf, Scl/Tal1 and Gata1 (*19*). Finally, the hematopoietic TF Scl/Tal1, a major actor in red blood cell differentiation (*19*), whose sequence-recognition binding motif is highly enriched in DMR sequences (Fig. S3F), is also expressed only in benthic fish (FDR<0.014; Fig. S5B), suggesting altered TF activity associated with methylome differences in benthic fish. Collectively, these results suggest significant divergence in epigenetic and transcriptional landscapes affecting erythropoiesis and haemoglobin composition in the benthic fish, which may facilitate occupation of anoxic conditions within the benthic habitat.

As noted above, the Lake Masoko populations are mainly characterised by an overall reduced transcriptional activity in many genes related to steroid metabolic pathways and energy homeostasis compared to the riverine fish (Fig. 2B). Many of these genes show significant gain in methylation compared to the riverine population. The examples we examined include the fatty acid desaturase 2 (fads2), part of the PPAR signalling pathway, with functions in fatty acid metabolism and dietary metabolic adaptations (*20*), as well as the catabolic enzyme inositol oxygenase (miox). Both genes show hypermethylation associated with weaker transcriptional activity in Masoko ecomorphs compared to the ancestral-related riverine fish (Fig. S7A-B). The interplay between metabolism and epigenetic variation has been well established (*8, 20, 21*), and suggests that epigenetic divergence might facilitate any differences in dietary resource use patterns during colonisation of Lake Masoko habitats.

### Plasticity and inheritance of population-specific methylome divergence

We examined the plasticity and inheritance of population-specific methylome divergence using a common garden experiment, whereby wild-caught *A. calliptera* specimens from Lake Masoko (both littoral and benthic populations), as well as from the adjacent river Mbaka, were bred and first-generation fish were reared under the same controlled laboratory conditions. Liver methylomes (WGBS) were then generated from two common-garden individuals for each population (Fig. S8A-B). Under a common rearing environment, and within one generation, the majority of DMRs (∼86%) found in wild populations of Lake Masoko were reset to resemble mostly unchanged methylome levels of riverine-ancestral fish (‘reset DMRs’), while the remaining ∼15% were retained/fixed between populations (‘fixed DMRs’; Fig. 3A-C and Figs. S8C and S9A), consistent with transgenerational retention of population-specific methylome patterns. Moreover, reset DMRs were on average almost twice as long as fixed DMRs (median: 424 and 227 bp, respectively; Fig. S9I). We then performed gene ontology enrichment analysis and found that reset DMRs were significantly enriched in the promoter of genes with functions related to liver metabolism, immune function and with DNA-binding activity (Fig. S9B-C; see also Fig. S4). Moreover, 77.4% of all differentially expressed genes found among the livers of wild populations were associated with reset methylome patterns upon environmental perturbation (Fig. S9C). Such genes include for example phosphatidate phosphatase LPIN2 involved in fatty acid metabolism, PRELI domain containing protein 3B (preli3b) implicated in phospholipid transport, and lanosterol 14-alpha demethylase (cyp51a1) with functions in sterol biosynthesis (Fig. 3D), suggesting a tight link between environmental conditions and methylome divergence associated with altered metabolism. Furthermore, the erythropoietic eklf and epoR, which have benthic-specific methylome and transcriptome patterns in wild fish, all resemble river and littoral highly methylated profiles in the common garden experiment (Fig. S9D).

**Fig. 3.**
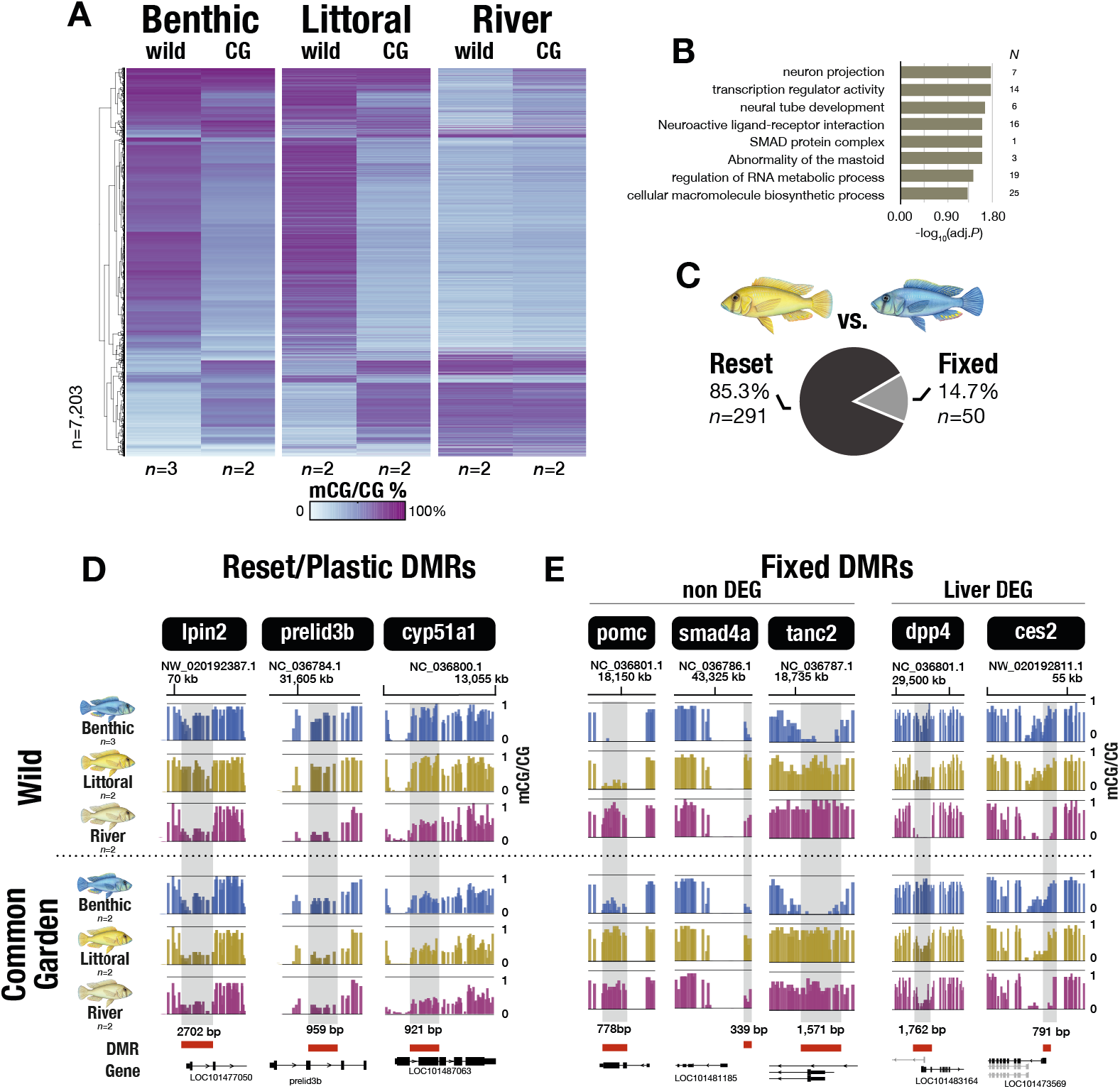
Common garden experiment results in global resetting of methylome profiles in wild benthic and littoral fish to resemble riverine methylome profiles, and inheritance of fixed methylome differences. **A.** Heatmap of the average DNA methylation levels (mCG/CG, %) at all DMRs found between wild populations in Fig. 1G for all wild and common-garden (CG) fish. Methylome profiles reveal global epigenetic resetting in wild benthic and littoral fish, to resemble the ancestral-like, riverine methylome profiles. **B**. Gene ontology enrichment analysis showing enrichment for fixed DMRs in developmental genes. **C**. The proportion of reset (upon common garden experiment and within one generation) and population-fixed DMRs between littoral and benthic fish. **D-E**. Examples of DMRs reset upon common garden experiment (D) and associated with population-specific transcriptional differences, and of DMRs fixed between populations (E) in wild and common-garden fish. Some fixed DMRs were also associated with altered transcriptional activity between populations (DEG, differentially expressed genes; see Fig. S9F-H). Each bar represents the average mCG/CG levels in 50bp-long non-overlapping windows for each fish population. DMRs (p<0.05) are highlighted in red and the length of each DMR is indicated in black.

While the majority of population-specific methylome patterns in wild populations show high levels of plasticity, suggesting a tight link between environmental conditions and epigenetic variation, methylome patterns fixed between populations show an overall significant association with genes related to development and embryogenesis, in particular brain development (Fig. 3B). Fixed DMRs might represent population-specific tissue-independent methylome patterns, possibly reflecting distinct core developmental processes (*7*). Examples of genes with functions during brain development and neuron morphogenesis and showing fixed population-specific methylome patterns include the embryonic transcription regulator SMAD family member 4a (smad4a; hypomethylated in littorals only) and the scaffolding protein tanc2 (hypomethylated in benthics) (Fig. 3E). Other pathways, related to biosynthetic processes among others, are also associated with fixed population-specific methylome patterns (Fig. 3B). This includes the gene pro-opiomelanocortin (pomc), a precursor polypeptide produced in the brain and involved in energy homeostasis and immune functions, showing retention of benthic-specific methylome patterns (Fig. 3E). Although three-quarters of all DEGs are associated with reset DMRs, we found some association between fixed population-specific methylome patterns and transcriptional divergence in liver-specific genes (22.6% of all DEGs), with functions enriched for catabolic and drug metabolism pathways (Fig. S10E). Two genes, in particular, coding for enzymes part of the insulin and fatty acid metabolic pathways respectively, dpp4 (dipeptidyl peptidase 4) and ces2 (liver carboxylase 2), show significant transcriptional downregulation in benthic livers and are associated with fixed benthic-specific hypermethylated levels at their promoters (Fig. 3E and Fig. S9G-H), suggesting retained epigenetic divergence associated with adaptation to different diets.

These results suggest that although methylome landscapes are highly environment-specific showing high plasticity and significantly associated with alteration of transcriptional activity of functional genes, some methylome divergence has become fixed and may be inherited in populations of Lake Masoko (fixed DMRs; Fig. 3D-E). While in mammals two waves in DNA methylation reprogramming occur early on, in zebrafish the paternal methylome is retained upon fertilisation, possibly allowing for transgenerational epigenetic inheritance (*22*) - however epigenetic reprogramming mechanisms in teleost fish may vary (*23*) and is currently unknown in cichlids.

## Conclusion

Our results provide direct evidence for functional and heritable methylome divergence associated with early stages of speciation in the radiation of *Astatotilapia* ecomorphs in Lake Masoko. Our analysis suggests that ancestral colonisation of the shallow lake habitat by a generalist riverine *Astatotilapia* population (∼10,000 years ago) was followed by colonisation of benthic habitat (∼1,000 years ago) characterised in part by both reversible and heritable methylome divergence in key functional genes (Fig. 4), including those related to haemoglobin synthesis, erythropoiesis, and sterol metabolism. We suggest that epigenetic processes provide the capacity for rapid occupation of new ecological niches prior to fixation of epigenetic and genomic variation. Our study therefore builds on evidence of epigenetic divergence seen among populations of fishes (*24*–*26*), intraspecific methylome remodelling in different rearing environments (*27, 28*), heritability of population-specific methylome divergence (*25, 29, 30*), and an epigenetic basis to diversification in vertebrate functional ecomorphological traits [eyes of cavefish *Astyanax mexicanus* (*4*)]. A key challenge now is to determine the mechanisms and rates of fixation of heritable epigenetic variation within populations (*6, 8, 22*), and how this associates with genomic fixation observed during the later stages of the speciation in cichlid fishes, and other model organisms used to study speciation and adaptive radiation (*4*–*6, 8, 9*).

**Fig. 4.**
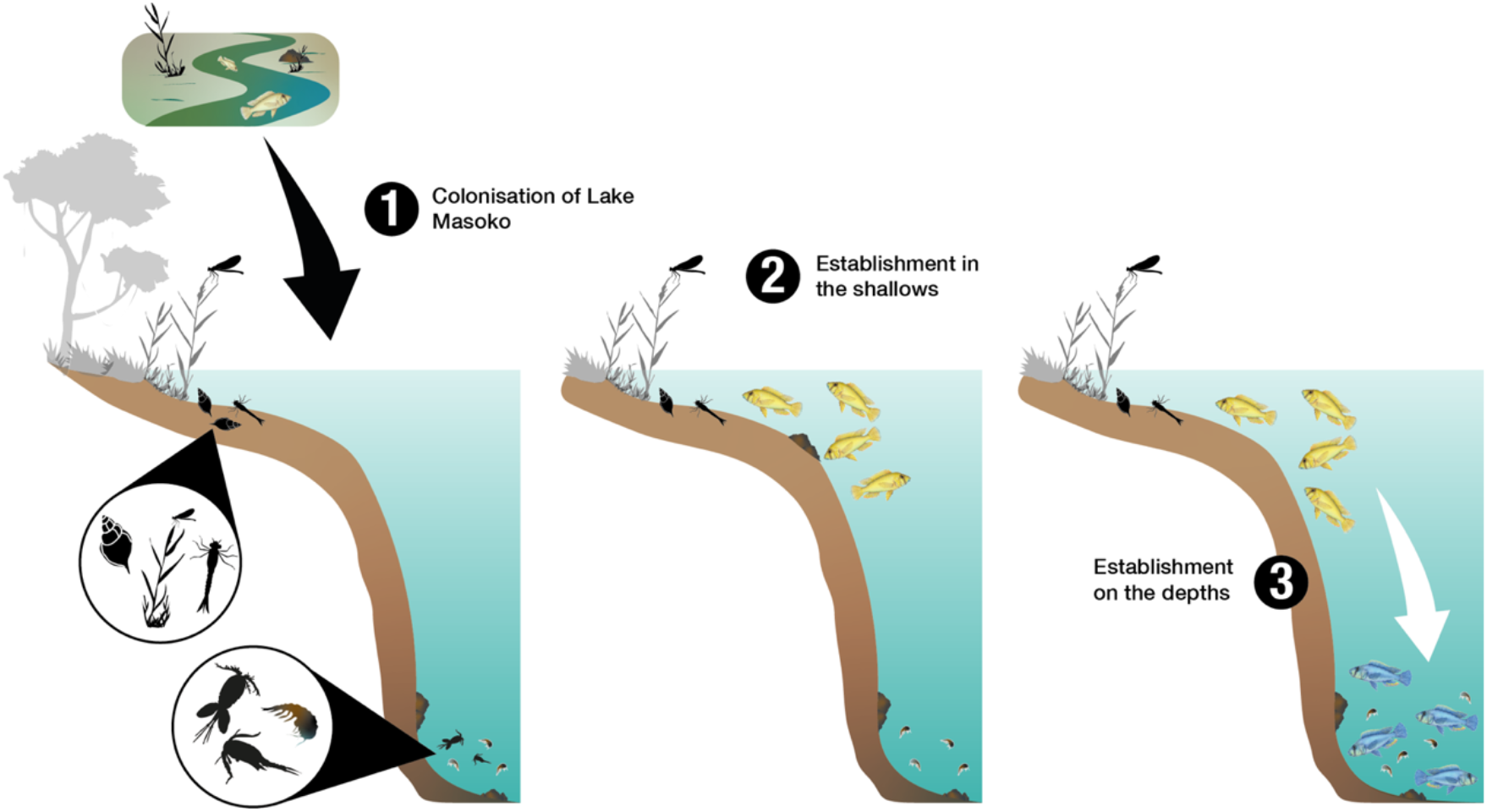
Hypothesised three stages of epigenetically-associated *Astatotilapia* ecomorph speciation in Lake Masoko. **1.** Ancestral colonisation of the shallow habitats of Lake Masoko by generalist riverine *Astatotilapia* population approximately 10,000 years ago (Malinsky et al., 2015). **2**. Occupancy of shallow, reedy highly oxygenated habitats by fish with a high level of depth philopatry. Phenotypic plasticity, partially linked to global methylome changes enables utilisation of littoral macroinvertebrate prey. **3**. Colonisation of deep, zooplankton-rich and low oxygen habitats by the shallow population ∼1,000 years ago (Malinsky et al., 2015). Extreme methylome changes in the benthic population associated with diet (e.g., fatty acid metabolism) and environment (e.g., haemoglobin composition). Epialleles become reciprocally fixed in the two populations, plausibly leading migrants, and those of intermediate epi-genotypes to suffer a fitness disadvantage. Eventually, selection leads to differential fixation of genomic variation.

## Supporting information

Supplementary Materials (including Supp Figures, and Materials/Methods)

## ACKNOWLEDGMENTS

We thank the staff of the Tanzania Fisheries Research Institute for their assistance and support. The Tanzania Commission for Science and Technology provided research permits, while the Tanzania Ministry of Livestock and Fisheries provided export permits. We thank Msafiri Ndawala (TAFIRI) and George Kazumbe for their help with sample collection, and Julie Johnson for her fish paintings. We thank Navin B. Ramakrishna for critical comments on the manuscript. for We thank the staff at the Gurdon Institute and the sequencing facilities at CRUK Cambridge Institute, Wellcome Gurdon and Wellcome Sanger Institutes, and Genomics facility at the University of Bristol for their expertise and support.

## AUTHOR CONTRIBUTION

<u>Sample collection</u>: GV, AGH, AS, BPN, NPG, MC, MES, AMT. <u>Common Garden experiment</u>: GFT. <u>RNA extraction</u>: GV, BF. <u>RRBS libraries and analysis</u>: AGH, GV. <u>WGBS libraries</u>: GV. <u>HDR annotation</u>, MM; <u>WGBS, RNAseq, RRBS analysis</u>: GV. <u>Supervision</u>: MJG, EAM, RD, GV, MES. <u>Conceptualisation</u>: MJG, GFT, EAM, GV. <u>Writing</u>: GV, MJG, EAM, with contribution from all the authors.

## DATA AND MATERIALS AVAILABILITY

WGBS, RRBS and RNA sequencing raw data have been deposited in the Gene Expression Omnibus (GEO) database and will be made public upon publication.

## SUPPLEMENTARY MATERIALS

Material and Methods

Supplementary Figures S1-S9

Supplementary Tables S1-S3

