## Supplementary Materials (including Supp Figures, and Materials/Methods) for "Epigenetic Divergence during Early Stages of Speciation in an African Crater Lake Cichlid Fish"

[1]Wellcome/CRUK Gurdon Institute, University of Cambridge, Cambridge CB2 1QN, UK;
[2]Department of Genetics, University of Cambridge, Cambridge CB2 3EH, UK;
[3]Wellcome Sanger Institute, Hinxton CB10 1SA, UK;
[4]School of Biological Sciences, University of Bristol, Bristol BS8 1QU, UK;
[5]Department of Zoology, University of Cambridge, Cambridge CB2 3EJ, UK;
[6]Tanzania Fisheries Research Institute (TAFIRI), Dar es Salaam, Tanzania;
[7]School of Natural Sciences, Bangor University, Bangor Gwynedd LL57 2UW, UK;
[8]Present address: School of Life Sciences, University of Hawai'i at Mānoa, Honolulu, USA;
[9]Present address: Max Planck Institute for Biology of Ageing, Cologne 50931, Germany;
[10]Present address: Institute of Ecology and Evolution, University of Bern, 3012 Bern, Switzerland.
[11]Equal contribution.

### MATERIALS AND METHODS

**Field sampling**

Lake Masoko fish were chased into fixed gill nets and SCUBA by a team of professional divers at different target depths determined by diver depth gauge (12x male benthic, 12x male littoral). Riverine fish (11x Mbaka River and 1x Itupi River) were collected by local fishermen. Upon collection, all fish were euthanised using clove oil. Collection of wild fish was done in accordance with local regulations and permits in 2015, 2016, 2018 and 2019. Upon collection, fish were immediately photographed with colour and metric scales and tissues were dissected and stored in RNA*later* (Sigma) - some samples were first stored in ethanol.

**Common garden experiment**

Common garden fish were bred from wild fish, collected and imported by a team of professional aquarium fish collectors in accordance with the veterinary regulations of the University of Bangor, UK. Fish were reared under the same controlled laboratory conditions in separate tanks (diet: algae flakes daily, 2-3times weekly frozen diet). Adult males (G1, generation 1) showing bright nuptial colours were culled using MS222 in accordance with veterinary regulations of the University of Bangor. Immediately upon culling, fish were photographed, and tissues collected and snap frozen in tubes.

**Isotope labelling**

Carbon ($\delta^{13}C$) and nitrogen ($\delta^{15}N$) isotope analysis of muscle samples (for same individuals as RRBS; 12, 12 and 9 samples for benthic, littoral riverine populations, respectively) was undertaken by Elemental Analysis - Isotope Ratio Mass Spectrometry (EA-IRMS) by Iso-Analytical Limited, UK. Dunn's multiple comparison tests following Kruskal-Wallis test were performed between the three groups using R package FSA v0.8.31 (p-values adjusted with the Holm method).

**RNAseq**

**NGS library prep**

Total RNA from liver tissues stored in RNA*later* was extracted using a phenol/chloroform approach (Trizol,Sigma). Quality and quantity of RNA extraction were assessed using TapeStation (Agilent), Qubit and NanoDrop (ThermoFisher). NGS libraries were prepared using polyA-tail-isolated RNA fraction and sequenced on NovaSeq (S4; paired-end 100/150bp), yielding on average 32.9±3.9 Mio reads.

**Read alignment and differential gene expression analysis**

Adaptor sequence in reads, low-quality bases (Phred score <20) and too short reads (<20bp) were removed using trimGalore (options: --paired --fastqc; v0.6.2; https://github.com/FelixKrueger/TrimGalore ). Paired-end reads were aligned to *M. zebra* reference genome (GCF_000238955.4_M_zebra_UMD2a_genomic.fa) using kallisto (*1*) (v0.46.0; options: --bias -b 100). Differential gene expression analysis was carried out with sleuth v0.30.0 (*2*) using Wald's test. Only genes with qval<0.05, fold change ≥1.5 between any pairwise comparison and showing high expression levels in ≥1 biological sample (maximal gene expression ≥10tpm in any one sample, which represents the 91$^{th}$ percentile for gene expression in the five benthic liver samples). Unbiased hierarchical clustering (complete-linkage clustering method) was carried out on R (3.6.3) using Euclidian's distances (dist) of Spearman's correlations (cor). Heatmaps were generated using pheatmap (v1.0.12; Euclidean distances and complete-linkage clustering method). Data for gene expression values (tpm) across different *A. calliptera* tissues were used from Vernaz et al., 2020.

**High-molecular-weight genomic DNA (HMW-gDNA) extraction**

HMW-gDNA from liver tissues stored in RNA*later* (Sigma) was isolated using QIAamp DNA Mini Kit (Qiagen 51304). Quality and quantity of extracted DNA samples were assessed using TapeStation (Agilent), Qubit and NanoDrop (ThermoFisher).

**Whole genome bisulfite sequencing (WGBS)**

**NGS library preparation (wild and common-garden samples)**

Unmethylated lambda phage genome (0.5% w/w) was first spiked in every sample (Promega D1521). DNA samples were then fragmented to ~400bp in length by sonication (Coveris E220). Length and quality of DNA fragments were assessed using TapeStation (Agilent). NGS libraries were prepared using ~400ng sonicated DNA fraction using NEBNext Ultra II DNA Library Kit for Illumina (E7645) and methylated adaptors (NEB E7535) according to manufacturer's instructions. DNA libraries were then treated with sodium bisulfite (Sigma Imprint, MOD50) according to manufacturer's instructions. Bisulfite-treated DNA libraries were then amplified by PCR (14cycles) and sequenced on Illumina HiSeq 4000 and NovaSeq (paired-end 150bp-long reads) to generate 322.02±58.94 million paired-end reads per sample (mean±sd).

**WGBS analysis**

Adaptor sequence in reads, low-quality bases (Phred score ≤20) and too short reads (<20bp) were removed using trimGalore (options: --paired --fastqc; v0.6.2; https://github.com/FelixKrueger/TrimGalore ). Sequencing reads (fastq) for the same sample generated on multiple lanes were merged. Paired-end reads were first mapped against lambda genome (GenBank accession: J02459) to assess bisulfite conversion (98.4±1.0%, mean±SD spike-in conversion rate), and to SNP-corrected version of *M. zebra* reference genome (GCF_000238955.4_M_zebra_UMD2a_genomic.fa) to account for *A. calliptera*-specific genotype/SNP (following same protocol developed in Ref.(*3*)) using Bismark v0.20.0 (options: -N 0 -p 4 -X 500) (*4*). Mapping rates were similar across samples, yielding 56.1±4.4% best unique reads mapping (mean±SD, n=13). Clonal paired-end reads (i.e., PCR duplicates) were removed using the deduplicate_bismark (options: -p --bam). Methylation scores (read count supporting mC/total read count) at each CpG site genome-wide were extracted using bismark_methylation_extractor (options: -p --multicore 6 --no_overlap --comprehensive --merge_non_CpG --bedGraph). Multiple HiSeq/NovaSeq lanes for one biological sample were merged. DMRs with ≥25% mCG difference, ≥4 CpG sites and p value<0.05 were predicted using DSS (*5*) (v2.34.0). DMRs predicted between wild Littoral and Benthic samples were considered fixed when found between common garden littoral and benthic fish (p<0.05) as well, or reset if not found (p>0.05). Wild DMRs found among wild riverine, littoral and benthic (Fig.3a) were merged when found in >1 pairwise comparison using bedtools (mergeBed).

For subsequent analyses, only CpG sites with ≥5 ≤100 unique (non-clonal) paired-end read coverage were used. Methylation scores at single CpGs were calculated using bismark output files as follows: number of methylated reads/total number of reads. Methylation levels in non-overlapping 50bp-long windows for each biological sample or each population (average mCG/CG) were generated with bedtools (v2.27.1; Ref (*6*)) and visualised as bigwig files (bedGraphToBigWig v.4, genome.ucsc.edu) in IGV genome browser (v.2.9.2; Broad Institute). Principal component analysis (centred and scaled) was carried out using R (v3.6.3; prcomp). Unbiased hierarchical clustering (complete-linkage clustering method) was carried out using R based on Euclidian's distances (dist) of pairwise Spearman's correlation scores (cor). Heatmaps were done using pheatmap (v1.0.12; Complete-linkage clustering method using Euclidean's distances). Circos plots were generated using circlise v0.4.12 to show DMR count across LG chromosomes only (NC chromosomes). Motif enrichment analysis within DMRs was performed on DMRs located outside gene bodies (excluding the first 1kbp downstream TSS) using HOMER v4.9 (findMotifs.pl to identify enriched motifs; scrambleFasta.pl on DMR fasta sequence to generate background sequence).

__Reduced representation bisulfite sequencing (RRBS)__

**NGS library prep and analysis**

HMW-gDNA from liver tissue from 12 adult male fish per ecomorph (36 total) was isolated using a modified version of the Wizard® Genomic DNA Purification Kit (Promega). Quality and quantity of extracted DNA samples were assessed using Qubit and NanoDrop (ThermoFisher). Approximatively 100 ng of liver HMW-gDNA were used to make RRBS libraries following the manufacturer's instructions (Premium RRBS kit, Diagenode, C02030032). Each of the three RRBS sequencing libraries multiplexed 12 different samples, with ecomorph representation randomized among libraries. Quality and quantity of all libraries were assessed using TapeStation (Agilent), Qubit and NanoDrop (ThermoFisher). RRBS libraries were sequenced on Illumina NextSeq500 (single-end 75bp long reads).

Due to poor read quality and low read counts, assessed using FastQC, one riverine ecomorph sample was excluded from further analysis. Analysis of spike-in controls gave a mean CpG bisulfite conversion efficiency across samples of 98.6%. Adaptor sequence in reads, low-quality end bases (Phred score ≤20), too short reads (<20bp) and the first 5bp (5'-end; to avoid sequencing bias) were removed using trimGalore v0.6.2 (options: --rrbs --fastqc --clip_R1 5; https://github.com/FelixKrueger/TrimGalore). In total after quality trimming, there were 11.1±3.4 Mio reads per RRBS sample (mean±sd). Reads were then aligned to *M. zebra* reference genome (GCF_000238955.4_M_zebra_UMD2a_genomic.fa) using Bismark v0.19.0 (options: -N 1; Ref. (*4*)). Mapping rates were 83.8±0.8%, 81.6±1.7% and 82.9±1.6%, for Benthic, River and Littoral populations, respectively. Methylation scores (read count supporting mC/total read count) at each CpG site genome-wide were extracted using bismark_methylation_extractor (options: -s --multicore 4 --comprehensive --merge_non_CpG --bedGraph). Principal component analysis of methylation levels at CpG sites found across all samples (common CG sites, n=151,900) was carried out using R prcomp(centered, scaled).

__Genomic annotation__

**DMR localisation**

Promoter regions are defined as TSS±1kbp. Gene bodies comprise exon and intron, minus the first 1kbp downstream of TSS. Only transposon-only repeats (TE) were analysed (excluding simple repeats, low complexity repeats, rRNA repeats and satellite repeats) - RepeatMasker annotation from Ref. (*3*). The annotation for CpG-islands (CGIs) was defined as in

Ref. (*3*). Intergenic regions were defined as regions outside promoters and gene bodies for DMR localisation. DMR coordinates were intersected with each respective genomic annotation and counted using bedtools.

**Enrichment for genomic features**

Enrichment for methylome divergence (DMR) in different genomic features was performed by dividing the observed number of DMR overlapping each genomic feature by the expected values (O/E ratio). The expected values were obtained by randomly shuffling the DMR coordinates genome-wide (1000x iterations) for each genomic feature. One sample t-tests were performed to test whether expected values were significantly different from the observed values. Chi-squared tests (R) were then performed for all O/E distributions among the three DMR comparison groups across all genomic features.

**DEG-DMR localisation**

DMRs were associated with DEG when located in either promoters (TSS±1kbp; Promoter DMRs), gene bodies (excl. the first 1kbp downstream TSS; Gene DMRs) or in the vicinity (1-5kbp away from closest gene; Intergenic DMRs) of the differentially expressed genes. Exact hypergeometric test (and representation factor) for the overlap between promoter-DMR and DEG was used.

**Gene Ontology Enrichment analysis**

All GO enrichments analyses were performed using g:Profiler (https://biit.cs.ut.ee/gprofiler/gost; version March 2021; (*7*)). Only annotated genes for Maylandia zebra were used with a statistical cut-off of FDR<0.05.

**Co-localisation with genetically Highly Diverged Regions (HDRs)**

The coordinates of HDRs from Malinsky et al. 2015 (*8*) were translated to the UMD2a *M. zebra* reference genome (GCF_000238955.4_M_zebra_UMD2a_genomic.fa) using the UCSC liftOver tool, based on a whole genome alignment between the original Brawand et al., 2014 (*9*) (https://www.ncbi.nlm.nih.gov/assembly/GCF_000238955.1) and the UMD2a M. zebra genome assemblies. The pairwise whole genome alignment was generated using lastz v1.02 (*10*), with the following parameters: "B=2 C=0 E=150 H=0 K=4500 L=3000 M=254 O=600 Q=human_chimp.v2.q T=2 Y=15000". This was followed by using USCS genome utilities (https://genome.ucsc.edu/util.html) axtChain tool with -minScore=5000. Additional tools with default parameters were then used following the UCSC whole-genome alignment paradigm (http://genomewiki.ucsc.edu/index.php/Whole_genome_alignment_howto) in order to obtain a contiguous pairwise alignment and the 'chain' file input for liftOver. All 98 HDRs mapped to the new assembly, although some HDR were split into more than one region in the UMD2a assembly, resulting in 141 regions. The distances between DMRs between littoral and benthic populations and the closest HDR were inferred using closestBed from the Bedtools2 suite (*6*).

### FIGURES S1 to S10

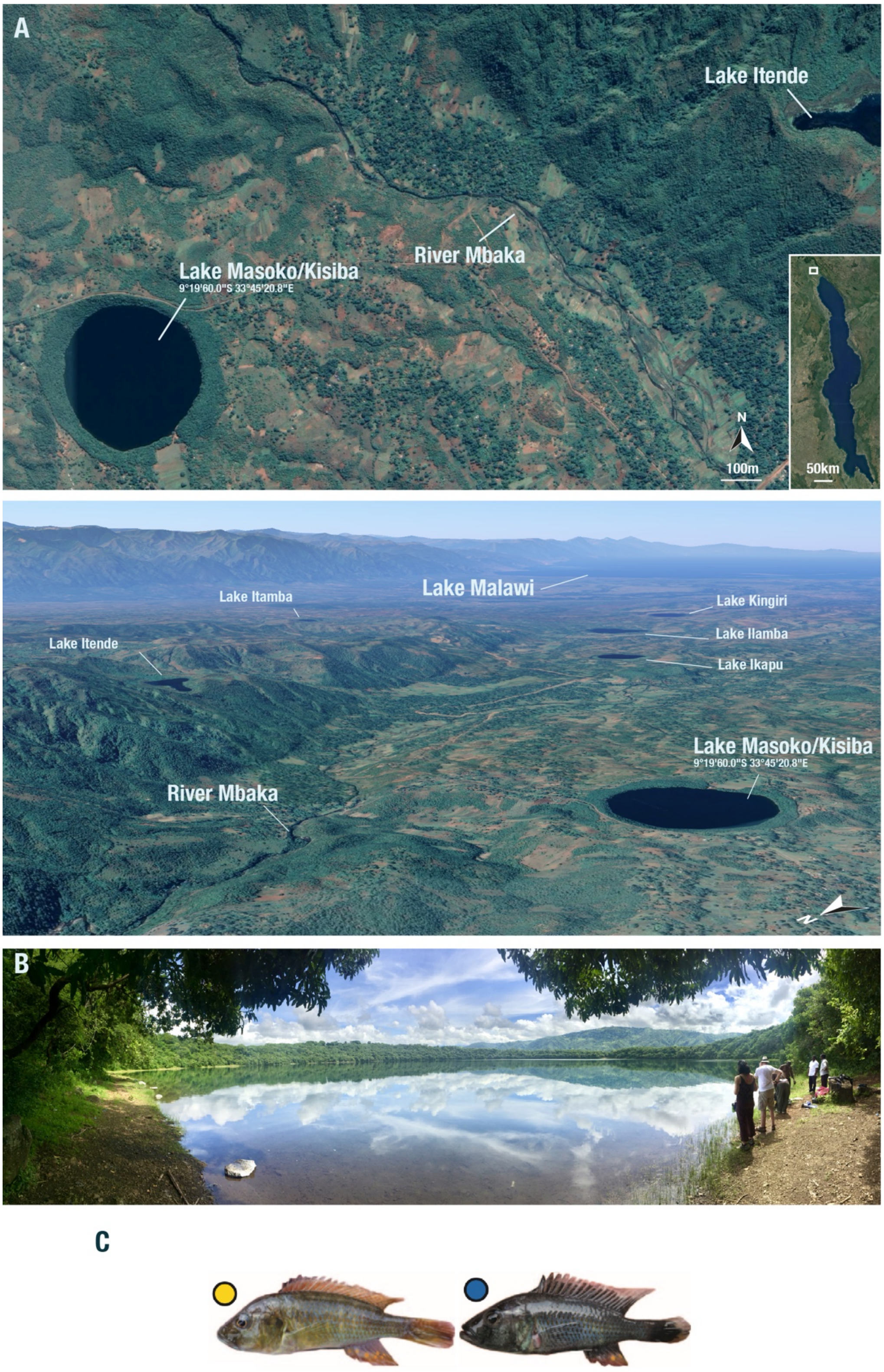

**Fig. S1. Map of Lake Masoko, Tanzania, and *Astatotilapia* cichlid ecomorphs of Lake Masoko.**

**A.** Upper: Satellite images of Lake Masoko/Kisiba and the neighbouring River Mbaka, Tanzania. Lower: Lakes and river of Lake Malawi catchment. Credits: Google Earth.

**B.** Panoramic photograph of Lake Masoko (11 April 2018) - photo credits: GV.

**C.** Photographs of male *A. calliptera* specimens of Lake Masoko in breeding colours: littoral ecomorph (left), benthic ecomorph (right). Photographs from Malinsky et al., Science. Ref. (*8*).

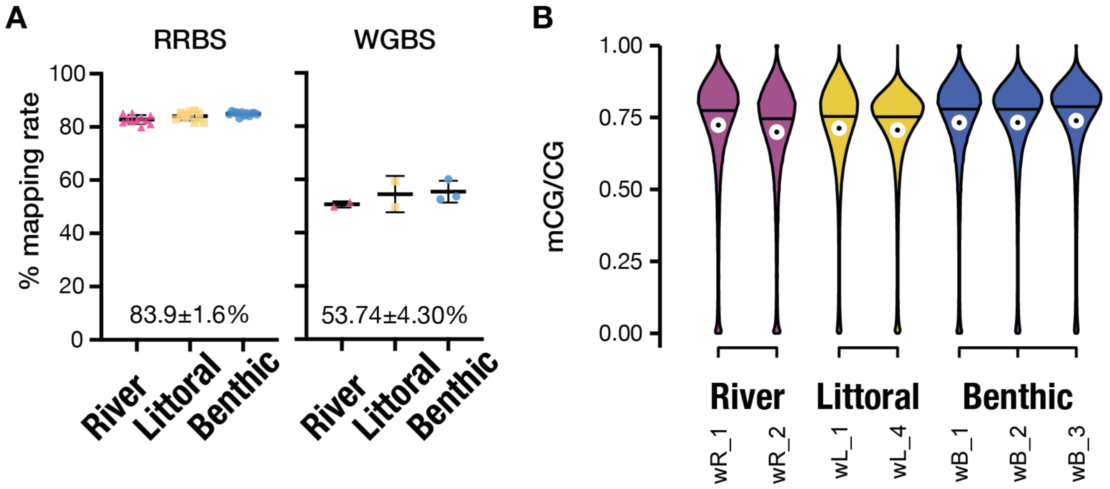

**Fig. S2. Mapping of sequencing reads and genome-wide methylome levels.**

**A.** Mapping rates for RRBS and WGBS reads aligned to the *Maylandia zebra* reference assembly (GCF_000238955.4 M_zebra_UMD2a). Black midlines and whiskers represent mean±sd of mapping rates for each population. Overall mapping rates are shown at the bottom of each graph (mean±sd).

**B.** Genome-wide liver methylation levels for each sample from each population (B, benthic, L, littoral and R, river; w, wild-caught). Average mCG/CG levels in non-overlapping 1kbp-long windows. n≥2 per population. Median and mean values are indicated with black midlines and white dots, respectively.

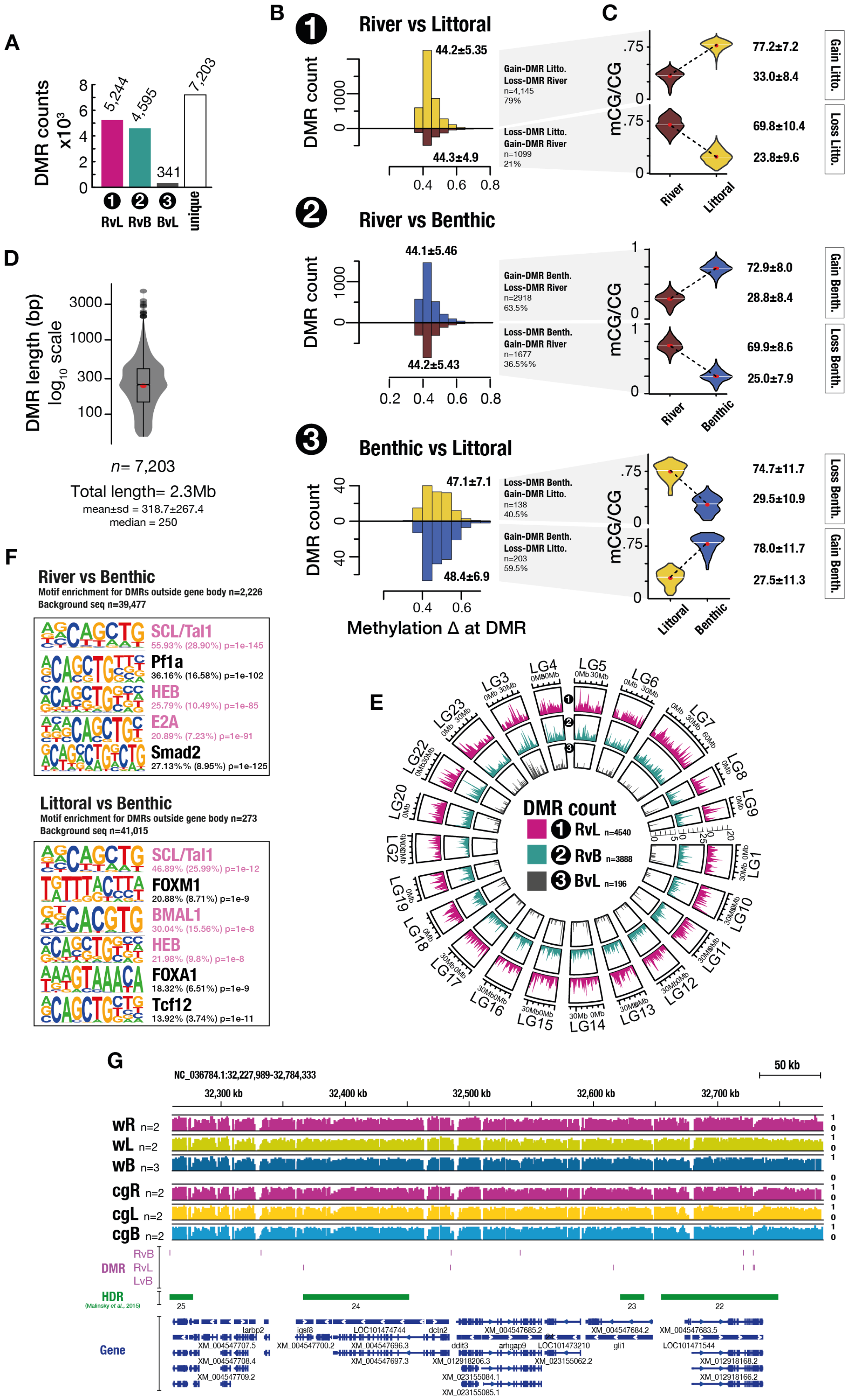

**Fig. S3. DMRs between wild populations of Lake Masoko and riverine *A. calliptera* fish.**

**A.** Total count of differentially methylated regions (DMR) found between each pairwise comparison (B, benthic, L, littoral and R, river; v., versus; see Fig. 1G) and total number of unique DMRs (found in ≥1 pairwise comparison; in grey).

**B.** Histograms of methylation difference (mCG/CG Δ) at DMRs found for each pairwise comparison. For each comparison, DMR are split between Gain/Loss DMRs for gain/loss of methylation. Gain-DMRs in benthic, littoral or river fish respectively

are indicated with histograms of different colours (blue, yellow or red, respectively). Average DMR methylation difference (mean±sd) is shown above/below each graph.
**C.** Violin plots showing average DMR mCG/CG levels for each DMR group found in (C). Values on the left of each graph represent mean±sd for mCG/CG levels.
**D.** Violin and box plots of the length (bp, log scale) of all unique DMRs found in ≥1 comparison. Mean value indicated by red point.
**E.** Circos plot showing DMR density found between each comparison (from Fig.1G) across all chromosomes (data shown only for linkage groups LG 1-23 in GCF_000238955.4 M_zebra_UMD2a).
**F.** Motif Enrichment analysis for transcription factor binding sites in DMR outside gene bodies over background (scramble DMR sequences), using HOMER (see Methods).
**G**. Methylome landscape (mean mCG/CG over 50bp windows) at four highly diverged regions (HDRs between littoral and benthic populations; from Malinsky et al., 2015) for wild (w) and common garden (cg) fishes from river (R), littoral (L) and benthic populations (n, biological replicates). DMR, differentially methylated regions in the three pairwise comparisons (v, versus). HDR annotations refer to Malinsky et al. 2015.

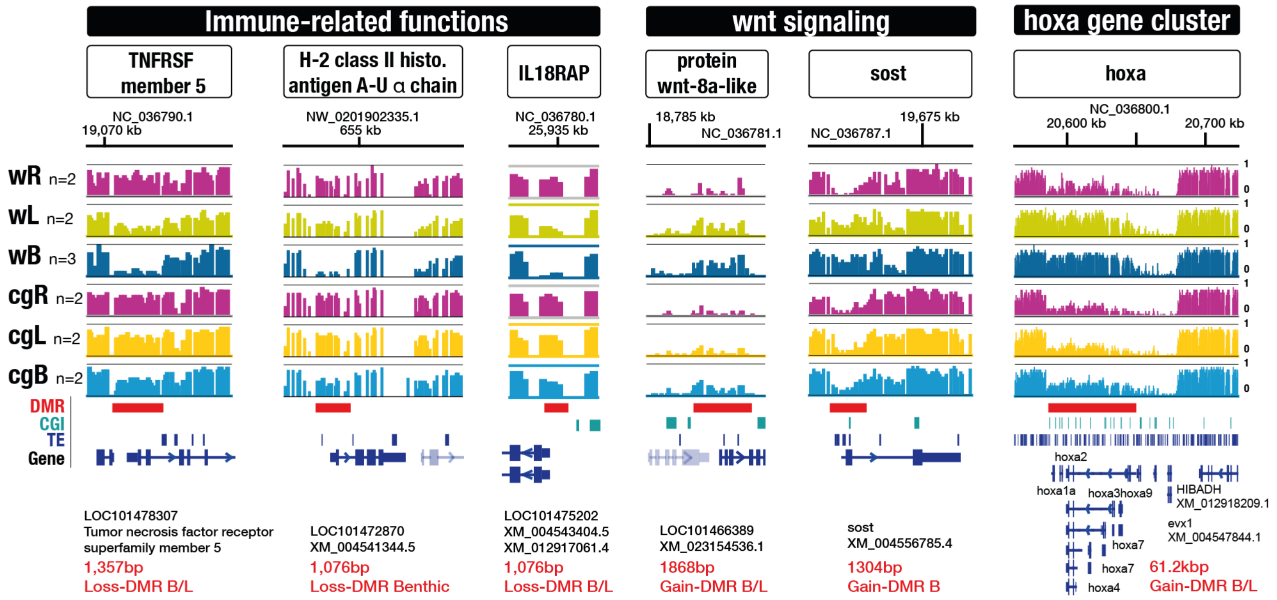

**Fig. S4. Population-specific DMRs are enriched in genes with functions related to immune system and development/embryogenesis.**

Examples of liver DMRs between the three populations (wild [w] and common garden [cg] fish). Average mCG/CG [0-1] in 50bp long windows (n indicate biological replicates). Lengths of DMRs are indicated in red at the bottom of each example. Loss-/Gain-DMRs indicate DMR showing significant decrease/increase in mCG/CG levels (at least 25% mCG/CG difference, $p<0.05$). **TNFRSF** member 5, tumor necrosis factor receptor superfamily member 5 (immune response, apoptosis); **H-2 class II histocompatibility antigen**, A-U alpha chain (adaptive immune response); **IL18RAP**, Interleukin-18 receptor accessory protein (immune response); **protein wnt-8a** and **sost**, sclerostin, both part of the wnt signalling pathway (early embryo embryogenesis). The homeobox **hoxa** genes cluster.

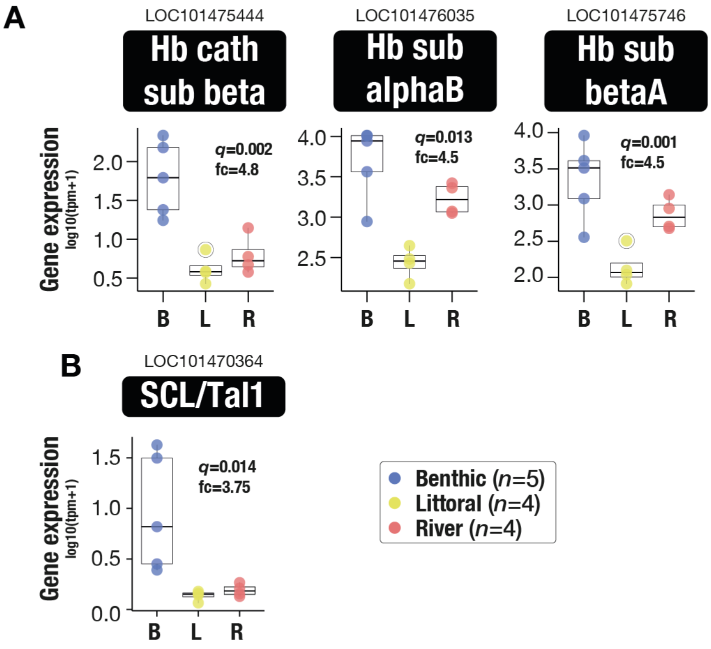

**Fig. S5. Genes with functions related to haematopoiesis and haemoglobin genes show high transcriptional activity in Benthic fish specifically.**

**A-B.** Boxplot of liver gene expression ($log_{10}$(tpm+1)) for the three haemoglobin (Hb) subunit genes (C) and for SCL/Tal1 gene (D), all significantly upregulated in benthic fish. *q*, false discovery rate adjusted p-value, using Benjamini-Hochberg (sleuth; Wald test) and fc ($log_2$(Fold change)) shown for Benthic vs Littoral comparison only.

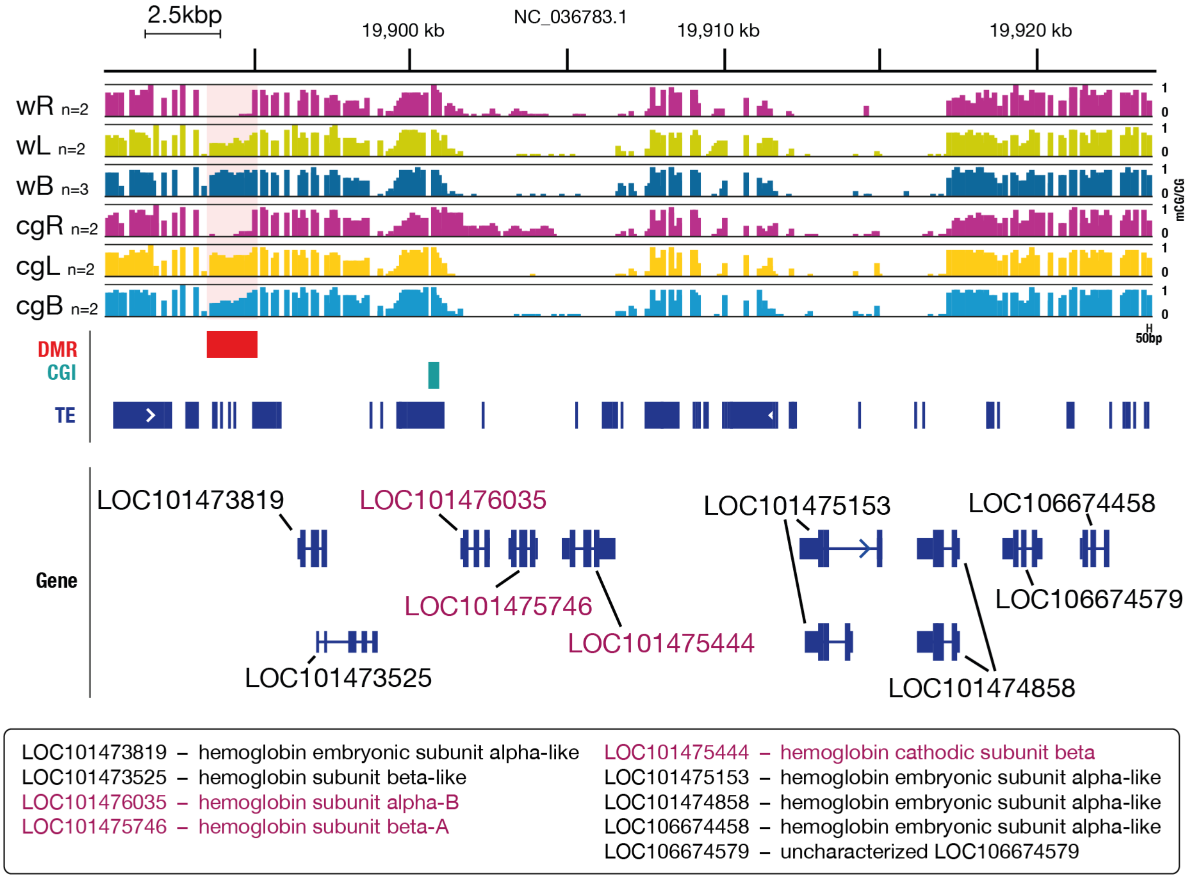

**Fig. S6. Large gain-DMR in the vicinity of the MN (major) globin cluster on chromosome LG4 in Benthic fish.** The 2kbp-long DMR is located 7kbp away from the MN globin cluster containing the three differentially expressed haemoglobin genes (see Supp Fig.S6c). Gain-DMR specifically in the Benthic fish. Littoral and river fish show intermediate and low methylation levels respectively. w, wild-caught; cg, common garden; R, L, B, river, littoral and benthic fish respectively. Each bar plot represents average mCG/CG levels for each population in 50bp-long windows.

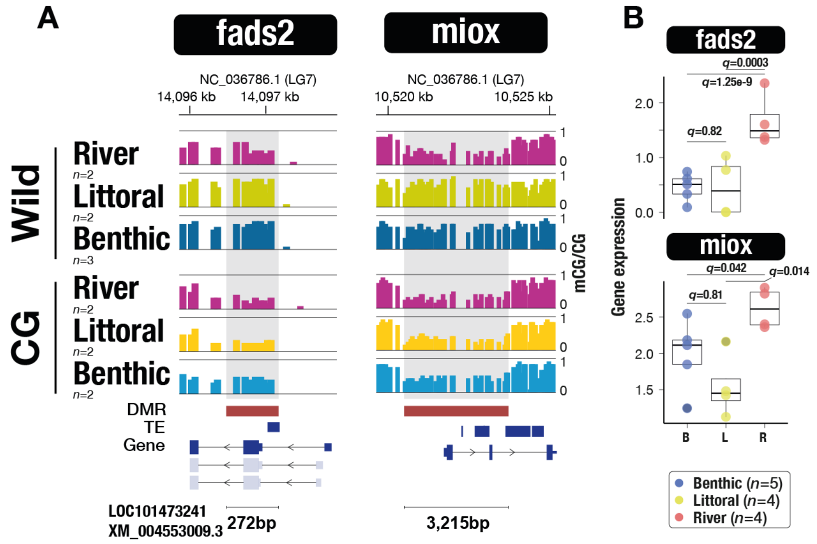

**Fig. S7. Gain-DMRs located in genes involved in fatty acid and steroid metabolic pathways and associated with altered transcriptional activity in the liver of both benthic/littoral fish compared to river fish.**

**A.** Methylation landscape in the genes fads2 (gene body) and miox (promoter). Each bar represents average liver mCG/CG level in 50bp-long windows for each population.

**B.** Boxplot of gene expression values (log10(tpm+1) for fads2 and miox in Benthic (n=5), Littoral (n=4) and River (n=4) liver tissues. qval shown for all comparisons.

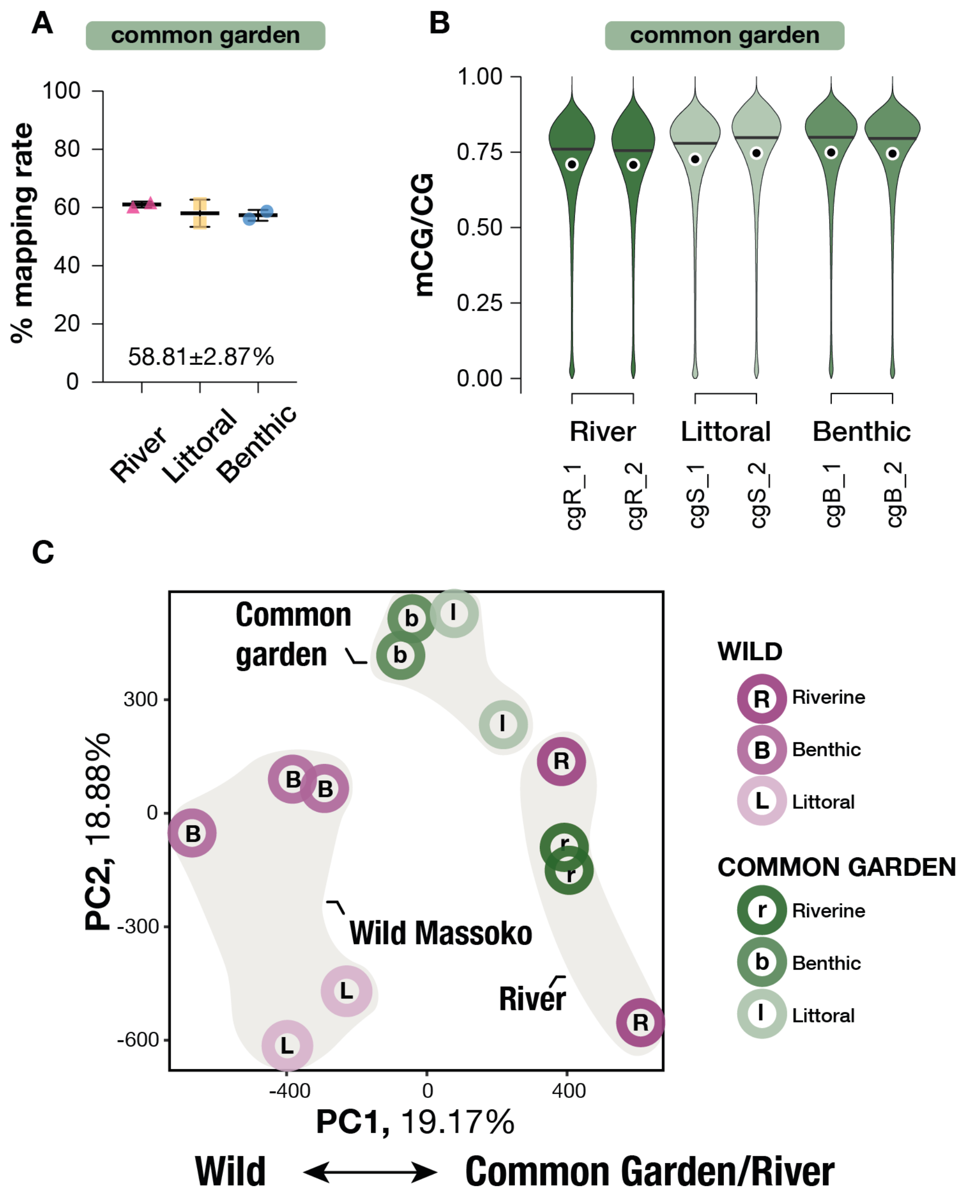

**Fig. S8. WGBS for Common-Garden fish and PC analysis.**

**A.** Mapping rates for Common Garden WGBS reads aligned to M.zebra_UMD2a reference genome. Black midlines and whiskers represent mean±sd of mapping rates within each population. Average mapping rates shown at the bottom of each graph (mean±sd).

**B.** Violin plots of average mCG/CG levels in 1kbp-long non-overlapping windows, genome wide, for wild and common garden fish in each population. cg, common garden fish; w, wild fish.

**C.** PCA of liver methylome variation in wild and common garden *A. calliptera* groups reveals global methylation remodelling in wild benthic and littoral fish (Wild Masoko) to resemble river fish methylome profiles. Wild and common garden fish specimens are highlighted in shades of purple and green colours, respectively. Upper- and lower-case letters distinguish wild from common garden fish for each group (river R/r, benthic B/b, littoral L/l).

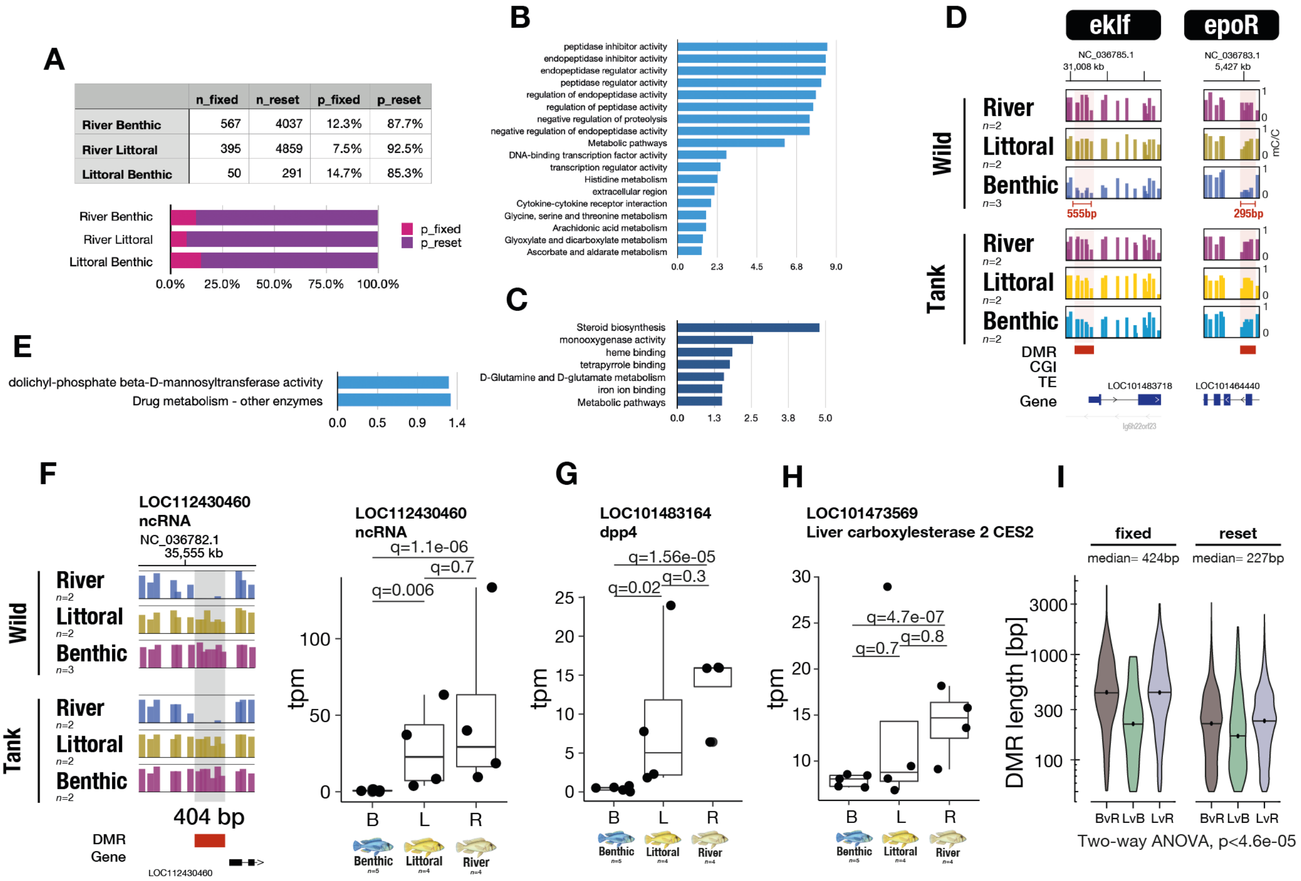

**Fig. S9. Fixed and reset methylome variation among wild populations of Lake Masoko *A. calliptera* cichlids.**

**A.** Plot showing the proportion of DMRs fixed vs reset between each pairwise comparison upon common garden experiment.

**B.** Gene ontology enrichment analysis using reset DMRs reveals enrichment for reset methylome patterns in genes with functions related to metabolism, immunity and with DNA binding activity.

**C.** GO analysis for fixed DMRs associated with differentially expressed genes.

**D.** Methylome profiles of the genes eklf and epoR for wild and common garden populations and showing reset methylome patterns in benthic fish upon environmental perturbation.

**E.** GO analysis for the differentially expressed genes associated with fixed methylome patterns.

**F.** Example of fixed benthic-specific hypomethylation in an uncharacterised non-coding RNA gene associated with its downregulation in benthic fish only.

**G-H.** Plot showing gene expression profiles for the genes dpp4 and ces2, associated with fixed benthic-specific methylome patterns.

**I.** Violin plots showing length in base pair (bp) of fixed and reset DMRs found between each pairwise comparison. y-axis, logarithmic scale. B, benthic; L, littoral; R, river populations respectively. Median length for all fixed and reset DMRs are indicated above the graphs. Two way ANOVA, *P*<4.6e05.

### TABLES S1-S4

**Table. S1. Sequencing summary for RRBS dataset.**

TableS1.txt

**Table. S2. Sequencing summary for WGBS dataset.**

TableS2.txt

**Table. S3. Sequencing summary for RNAseq dataset.**

TableS3.txt
